## Supplemental Table 2 for "Association of genetic variation in *COL11A1* with adolescent idiopathic scoliosis"

| **VarID_GRCh37** | **RsID** | **Mutation** | **gnomAD** | **CADD** | **GERP** | **#Cases** |
| --- | --- | --- | --- | --- | --- | --- |
| 1:103377744:C:T | rs151249006 | NM_080629.2:c.G4093A:p.A1365T | 6.37E-05 | 25.8 | 5.42 | 1 |
| 1:103381192:C:A | rs150669855 | NM_080629.2:c.G3847T:p.V1283L | 0.0007 | 0.001 | -10.9 | 4 |
| 1:103405909:C:T | rs370589018 | NM_080629.2:c.G3394A:p.G1132S | 6.37E-05 | 25.2 | 5.46 | 1 |

Supplemental Table 2. Rare *COL11A1* variants detected in 625 AIS exomes.
