## Supplemental Figures for "Association of genetic variation in *COL11A1* with adolescent idiopathic scoliosis"

### Slide 1
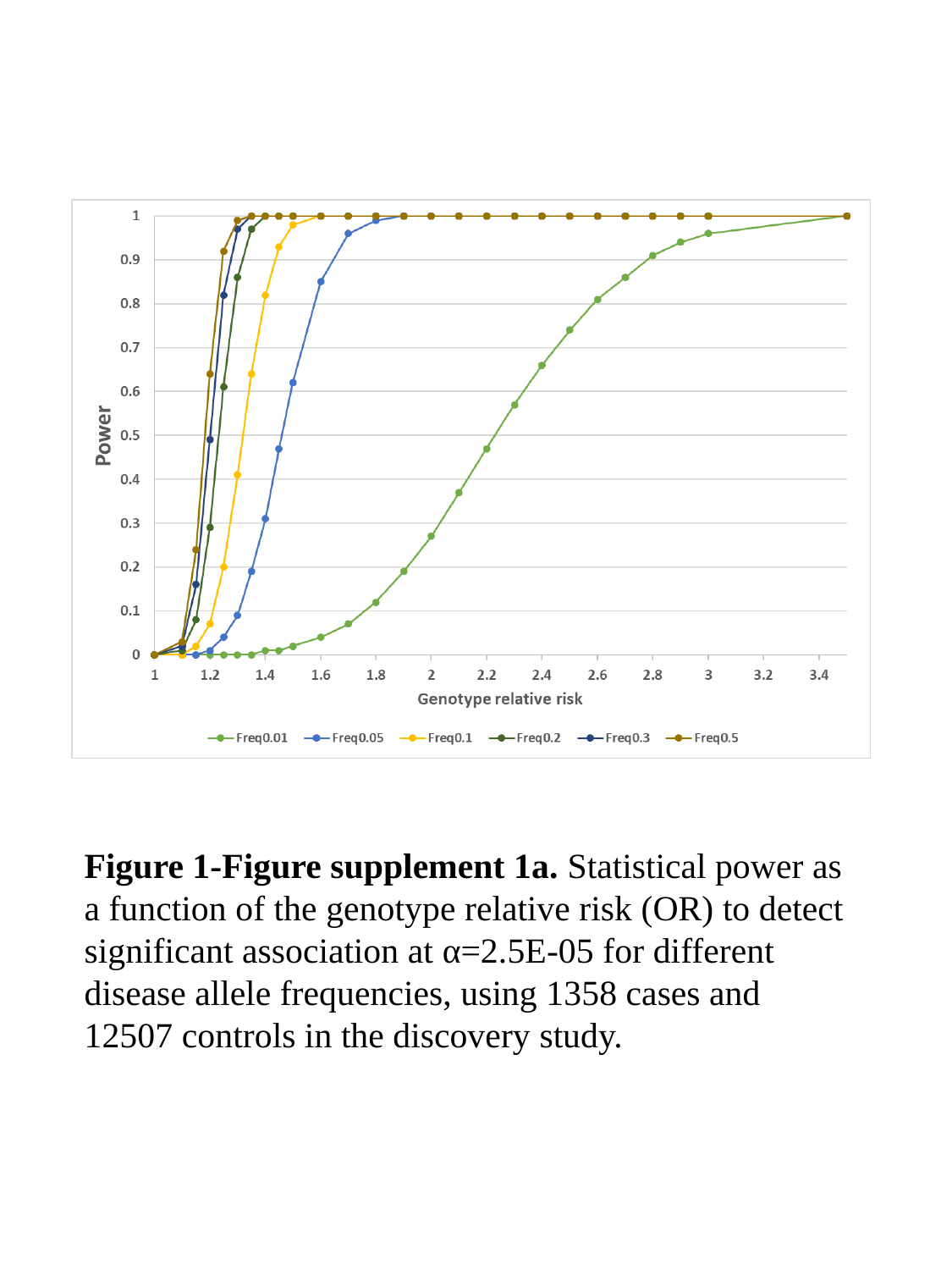

Figure 1-Figure supplement 1a. Statistical power as a function of the genotype relative risk (OR) to detect significant association at α=2.5E-05 for different disease allele frequencies, using 1358 cases and 12507 controls in the discovery study.

### Slide 2
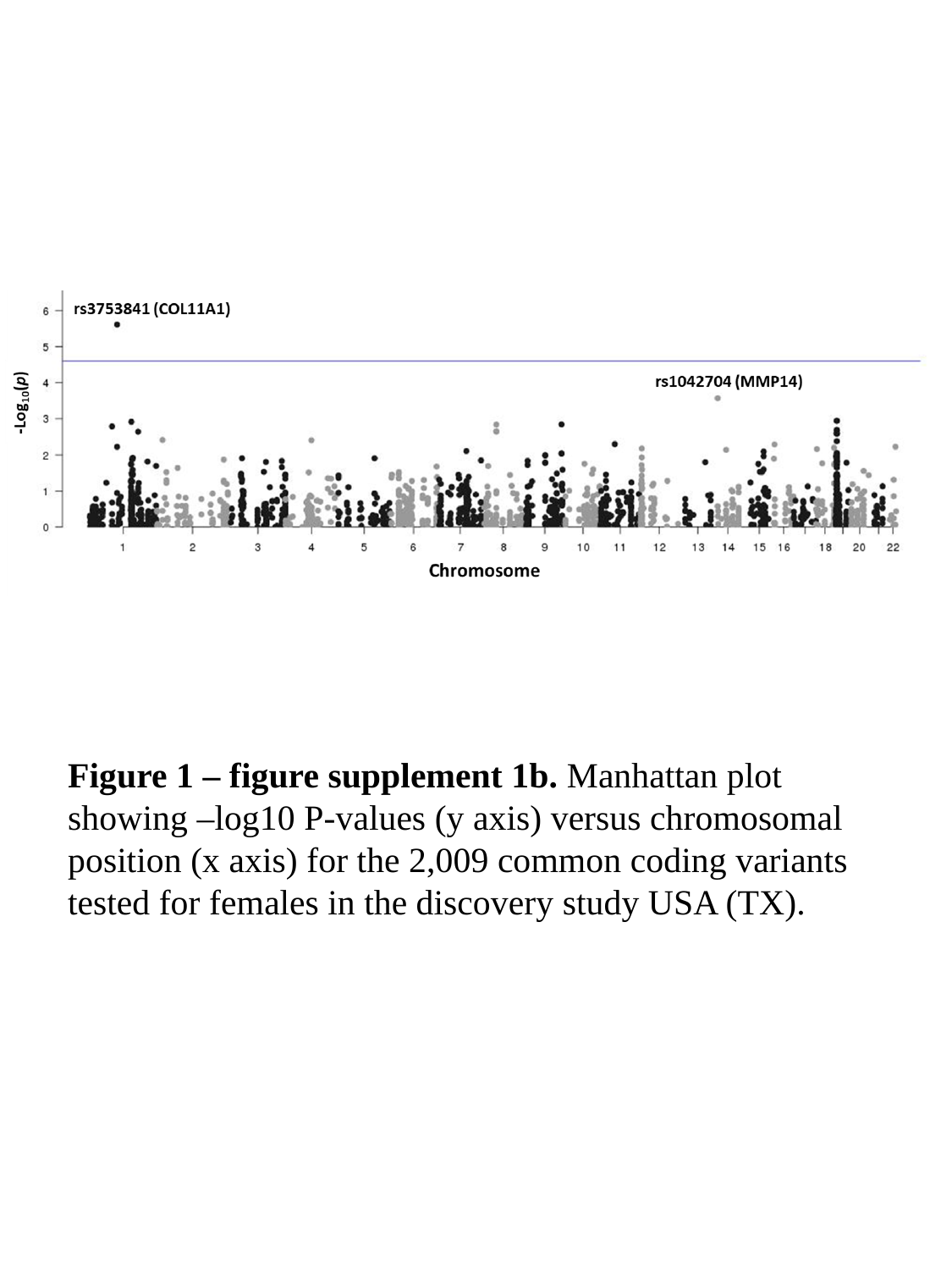

Figure 1 – figure supplement 1b. Manhattan plot showing –log10 P-values (y axis) versus chromosomal position (x axis) for the 2,009 common coding variants tested for females in the discovery study USA (TX).

### Slide 3
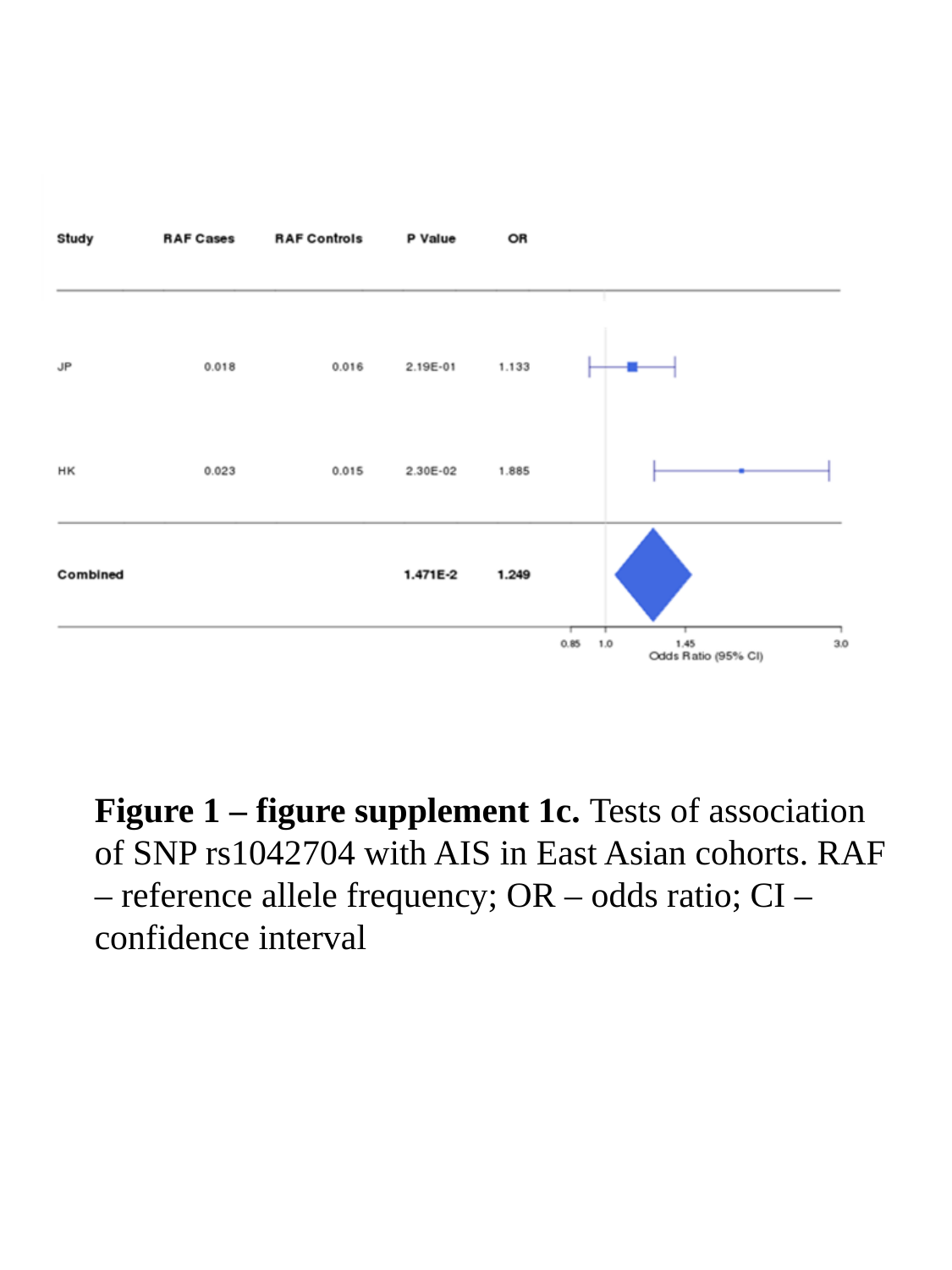

Figure 1 – figure supplement 1c. Tests of association of SNP rs1042704 with AIS in East Asian cohorts. RAF – reference allele frequency; OR – odds ratio; CI – confidence interval

### Slide 4
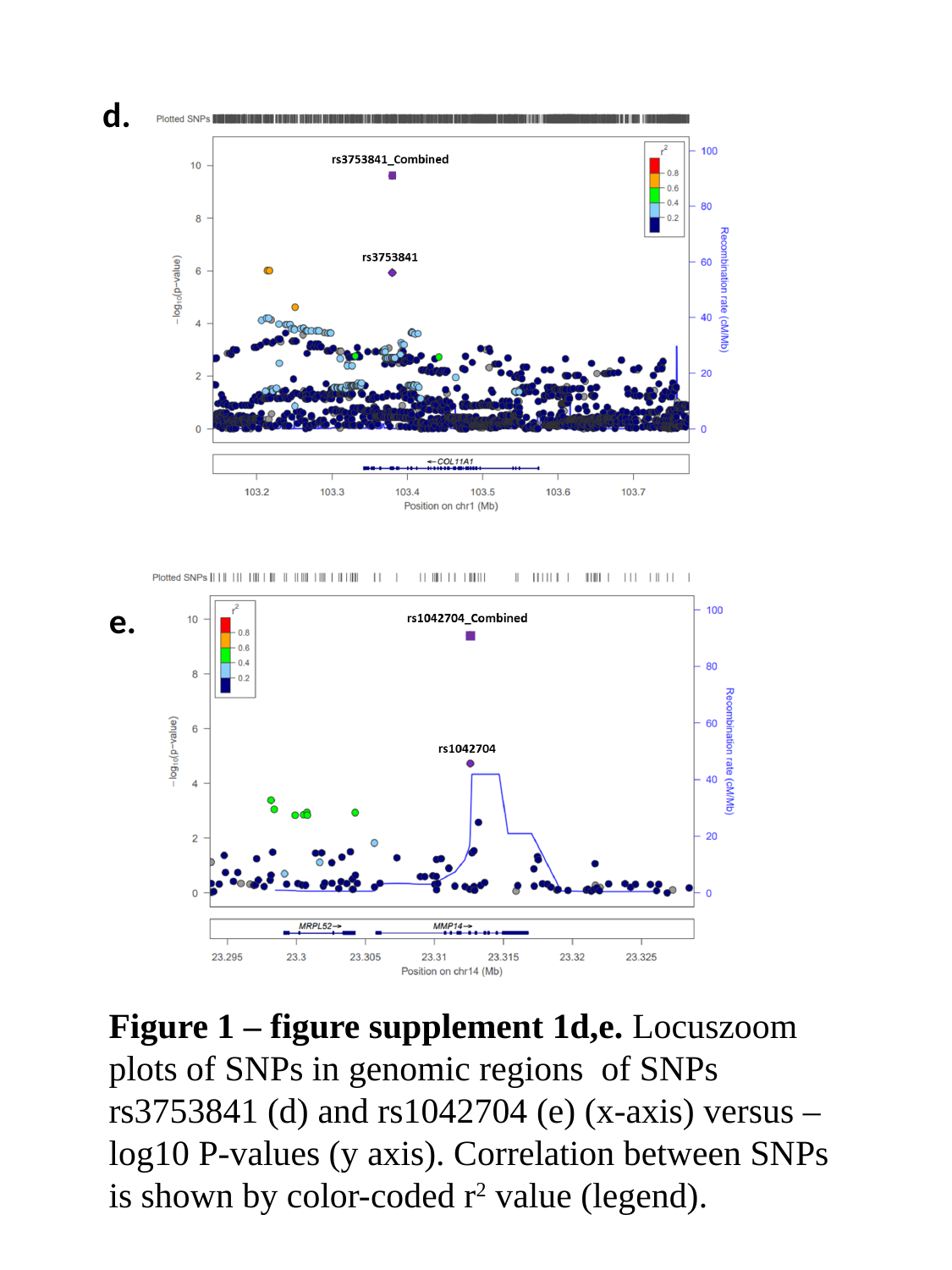

d.
e.
Figure 1 – figure supplement 1d,e. Locuszoom plots of SNPs in genomic regions of SNPs rs3753841 (d) and rs1042704 (e) (x-axis) versus –log10 P-values (y axis). Correlation between SNPs is shown by color-coded r^2^ value (legend).

### Slide 5
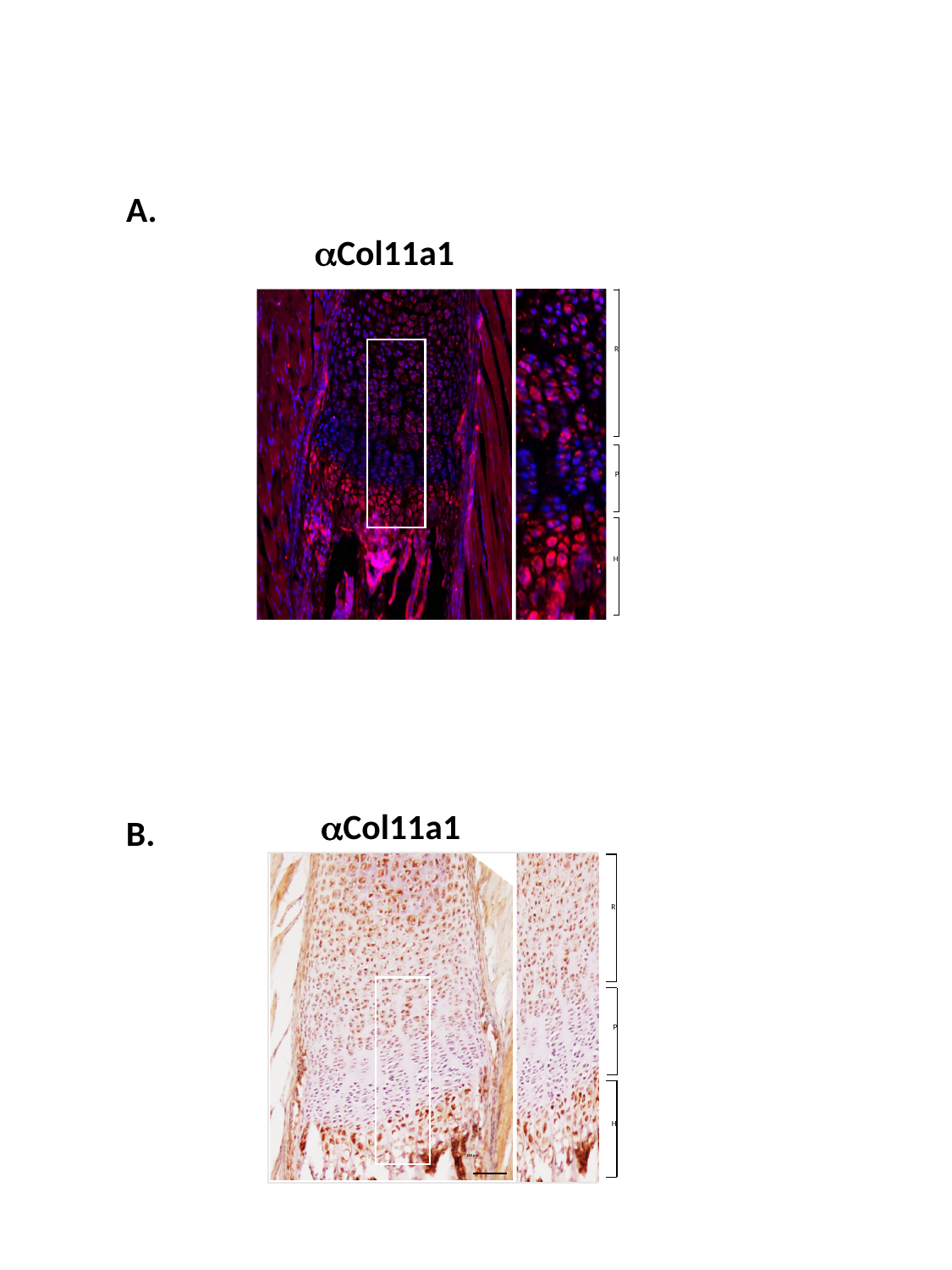

A.
αCol11a1
R
P
H
100 µm
αCol11a1
B.
R
P
H
100 µm

### Slide 6
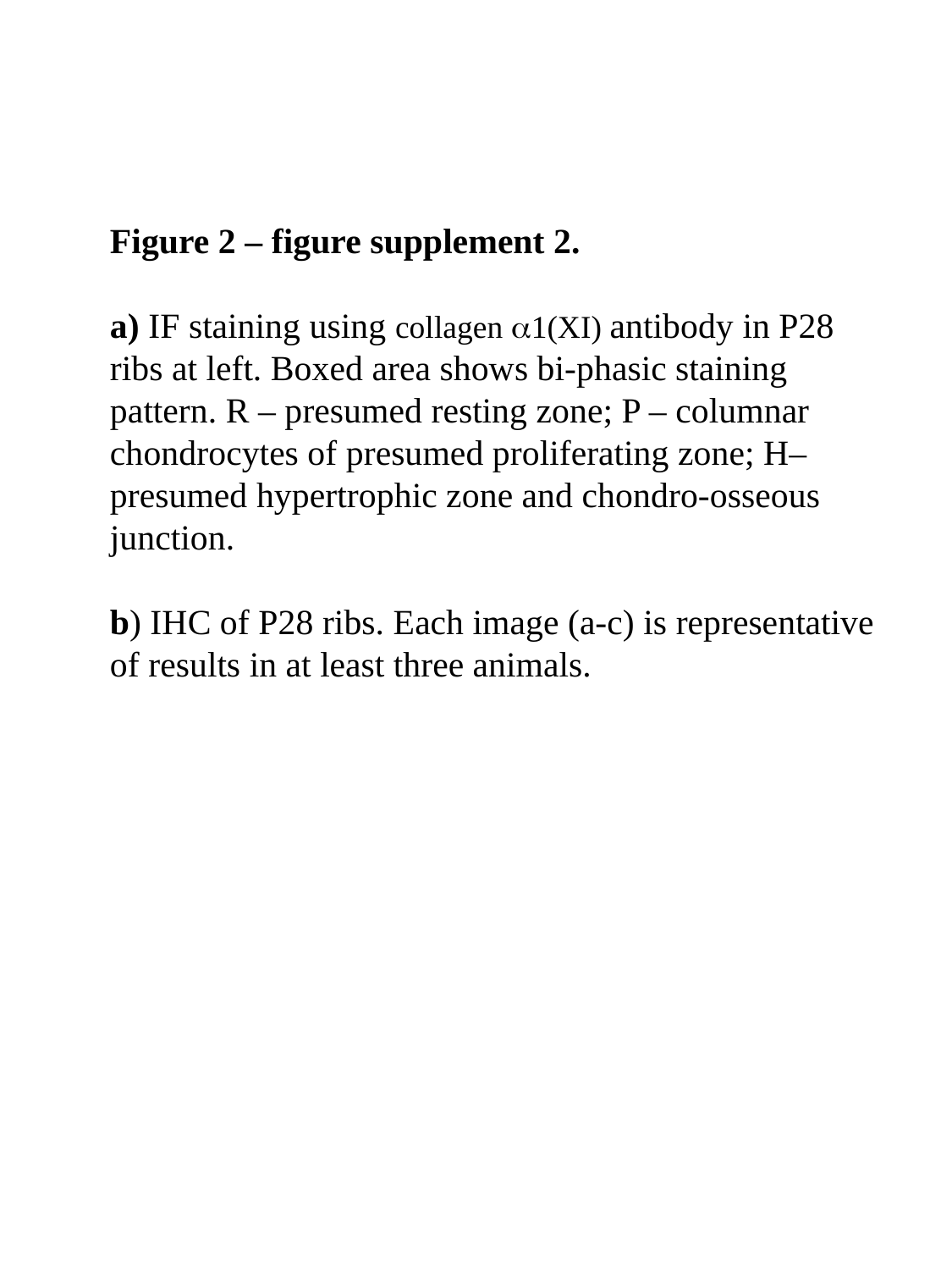

Figure 2 – figure supplement 2.
a) IF staining using collagen α1(XI) antibody in P28 ribs at left. Boxed area shows bi-phasic staining pattern. R – presumed resting zone; P – columnar chondrocytes of presumed proliferating zone; H– presumed hypertrophic zone and chondro-osseous junction.
b) IHC of P28 ribs. Each image (a-c) is representative of results in at least three animals.

### Slide 7
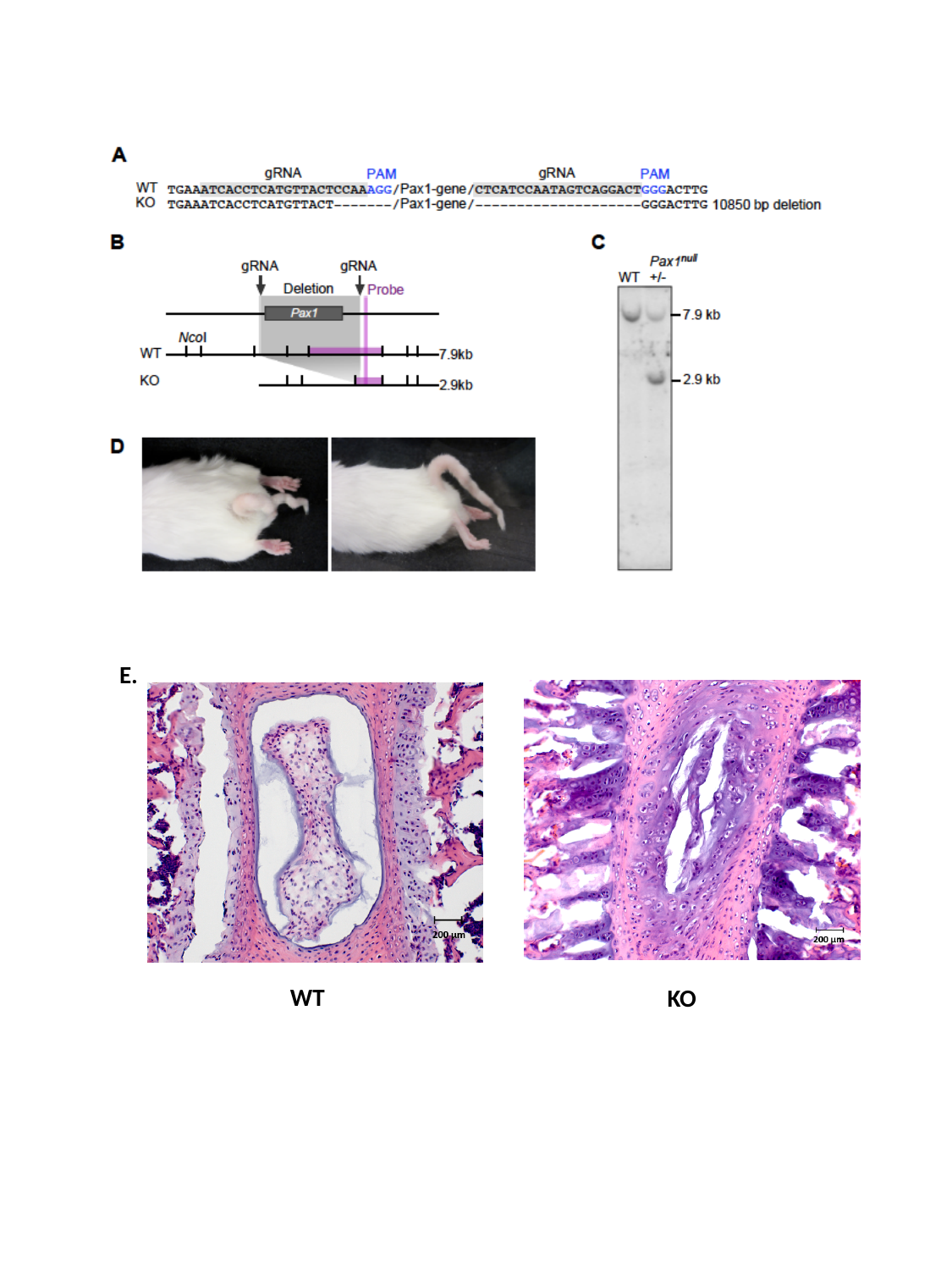

E.
WT
KO

### Slide 8
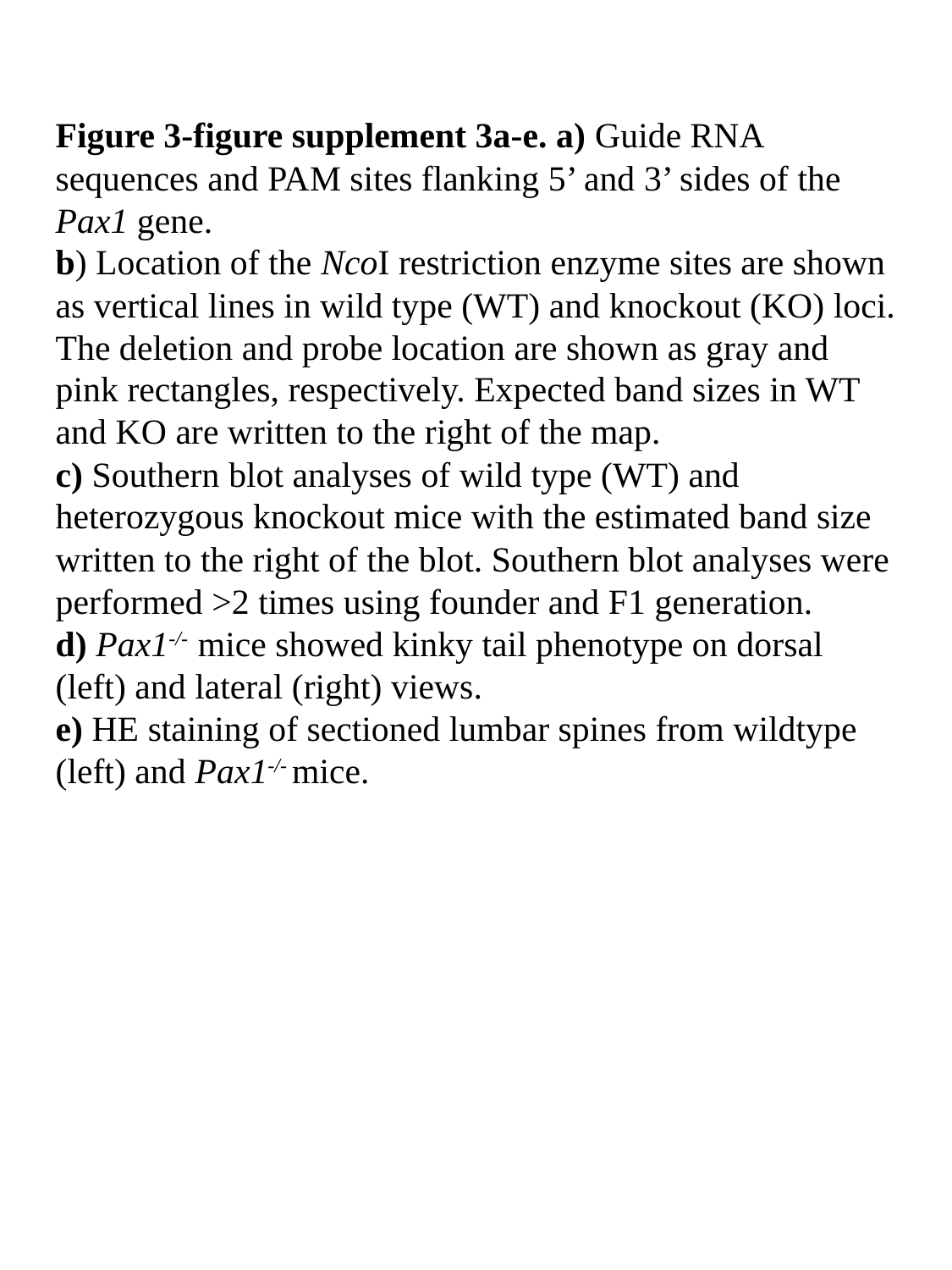

Figure 3-figure supplement 3a-e. a) Guide RNA sequences and PAM sites flanking 5’ and 3’ sides of the Pax1 gene.
b) Location of the NcoI restriction enzyme sites are shown as vertical lines in wild type (WT) and knockout (KO) loci. The deletion and probe location are shown as gray and pink rectangles, respectively. Expected band sizes in WT and KO are written to the right of the map.
c) Southern blot analyses of wild type (WT) and heterozygous knockout mice with the estimated band size written to the right of the blot. Southern blot analyses were performed >2 times using founder and F1 generation.
d) Pax1^-/-^ mice showed kinky tail phenotype on dorsal (left) and lateral (right) views.
e) HE staining of sectioned lumbar spines from wildtype (left) and Pax1^-/-^ mice.

### Slide 9
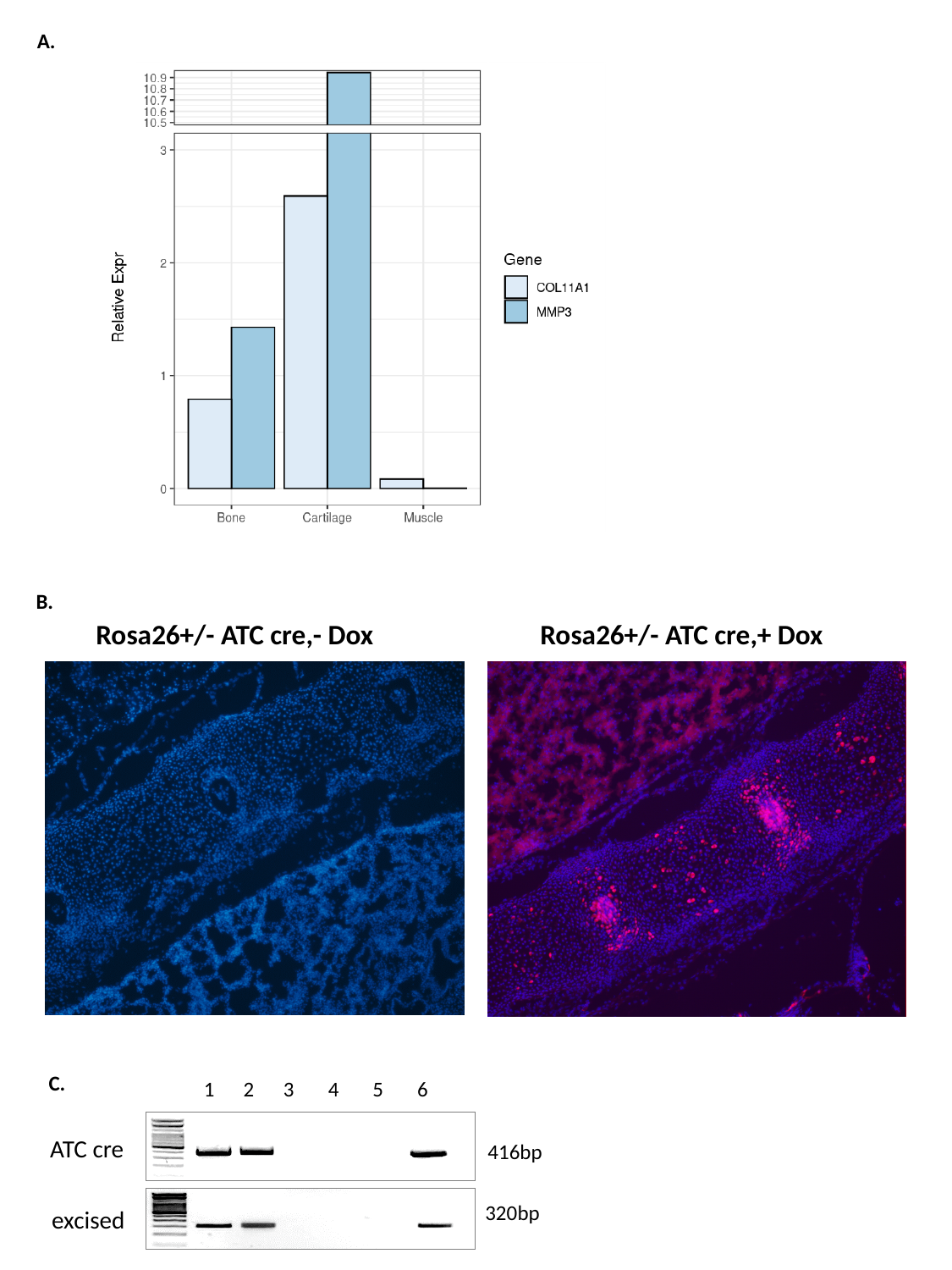

A.
B.
Rosa26+/- ATC cre,- Dox
Rosa26+/- ATC cre,+ Dox
C.
1 2 3 4 5 6
ATC cre
416bp
320bp
excised

### Slide 10
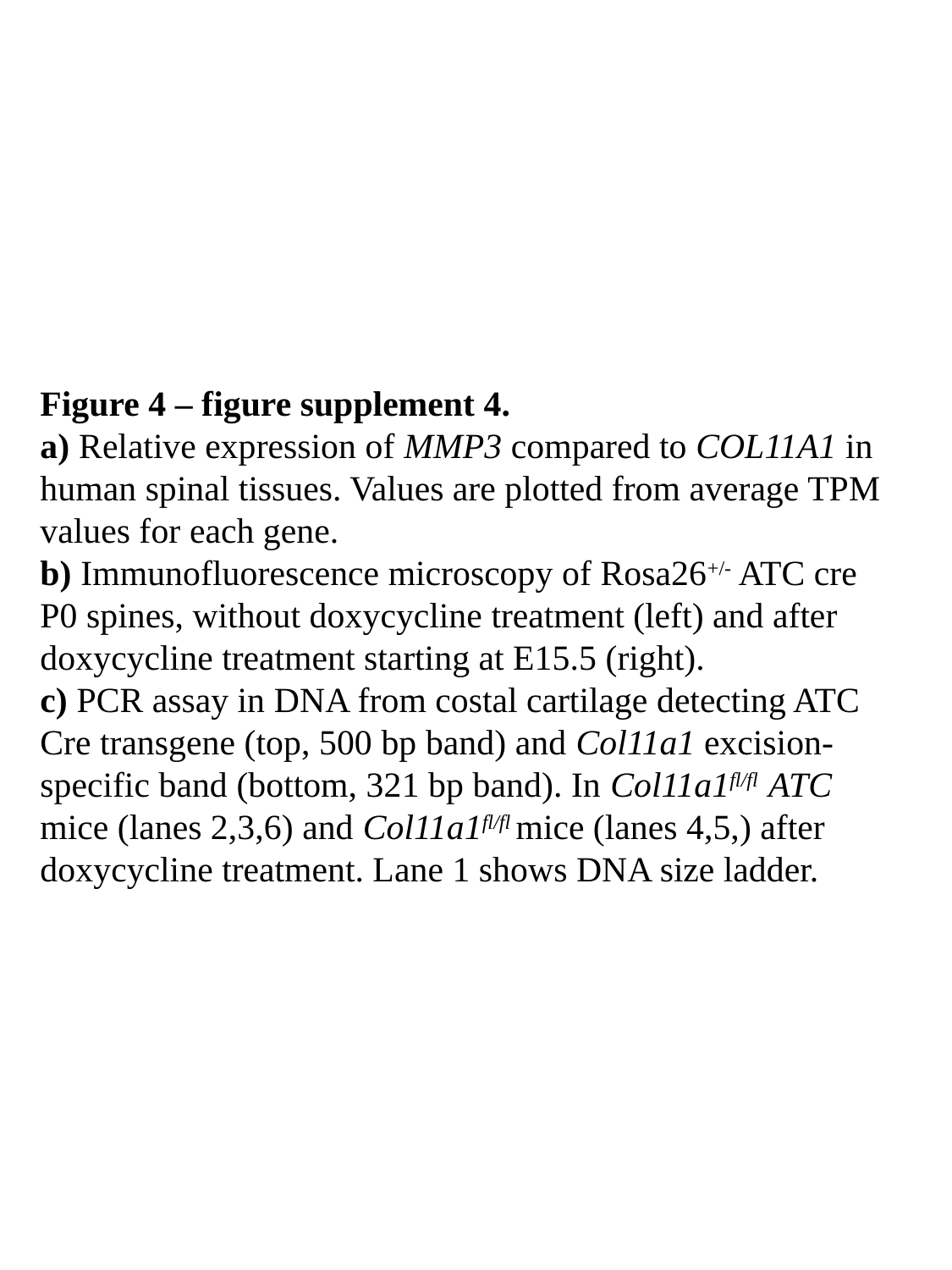

Figure 4 – figure supplement 4.
a) Relative expression of MMP3 compared to COL11A1 in human spinal tissues. Values are plotted from average TPM values for each gene.
b) Immunofluorescence microscopy of Rosa26^+/-^ ATC cre P0 spines, without doxycycline treatment (left) and after doxycycline treatment starting at E15.5 (right).
c) PCR assay in DNA from costal cartilage detecting ATC Cre transgene (top, 500 bp band) and Col11a1 excision-specific band (bottom, 321 bp band). In Col11a1^fl/fl^ ATC mice (lanes 2,3,6) and Col11a1^fl/fl^ mice (lanes 4,5,) after doxycycline treatment. Lane 1 shows DNA size ladder.

### Slide 11
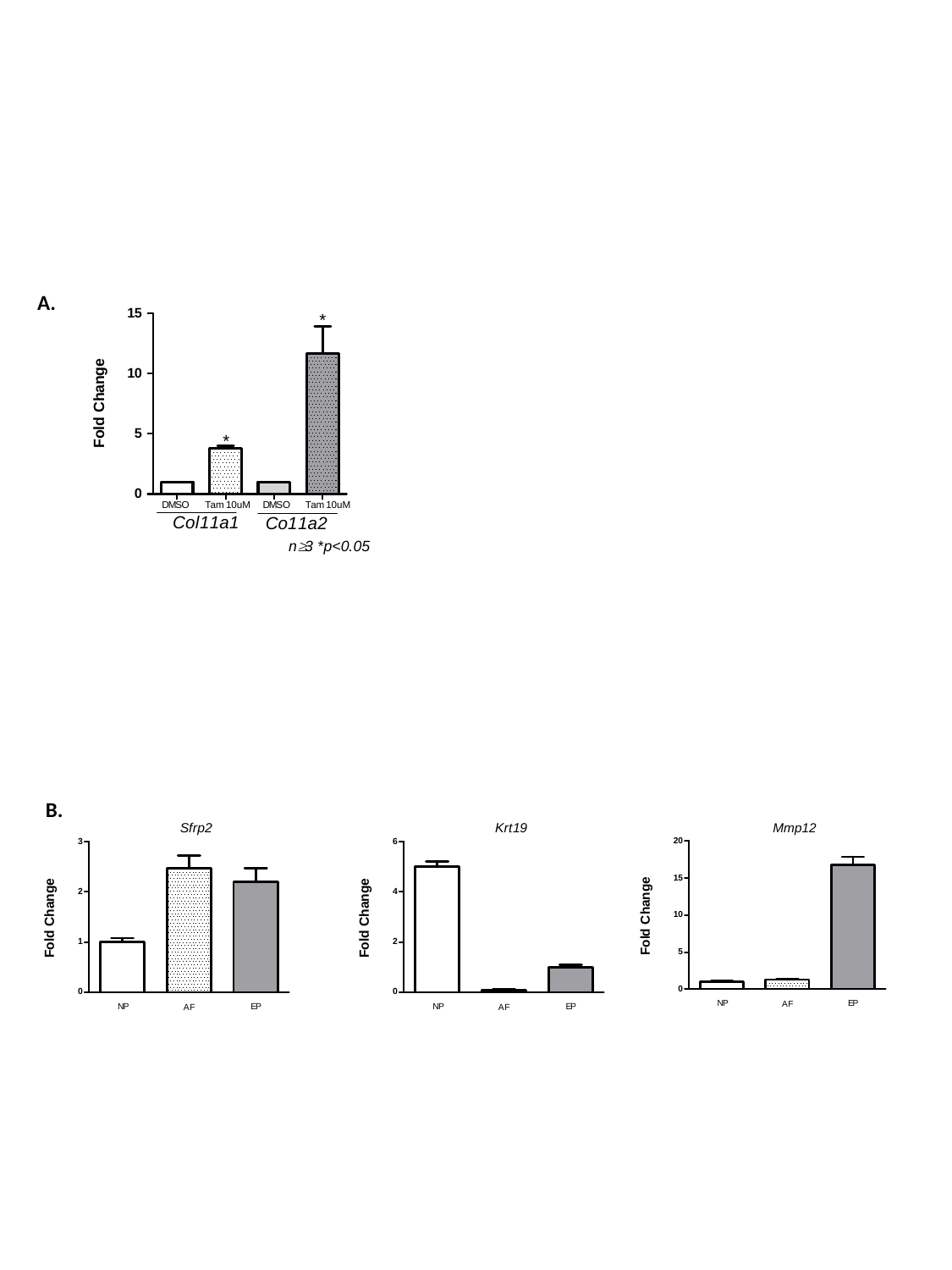

A.
B.

### Slide 12
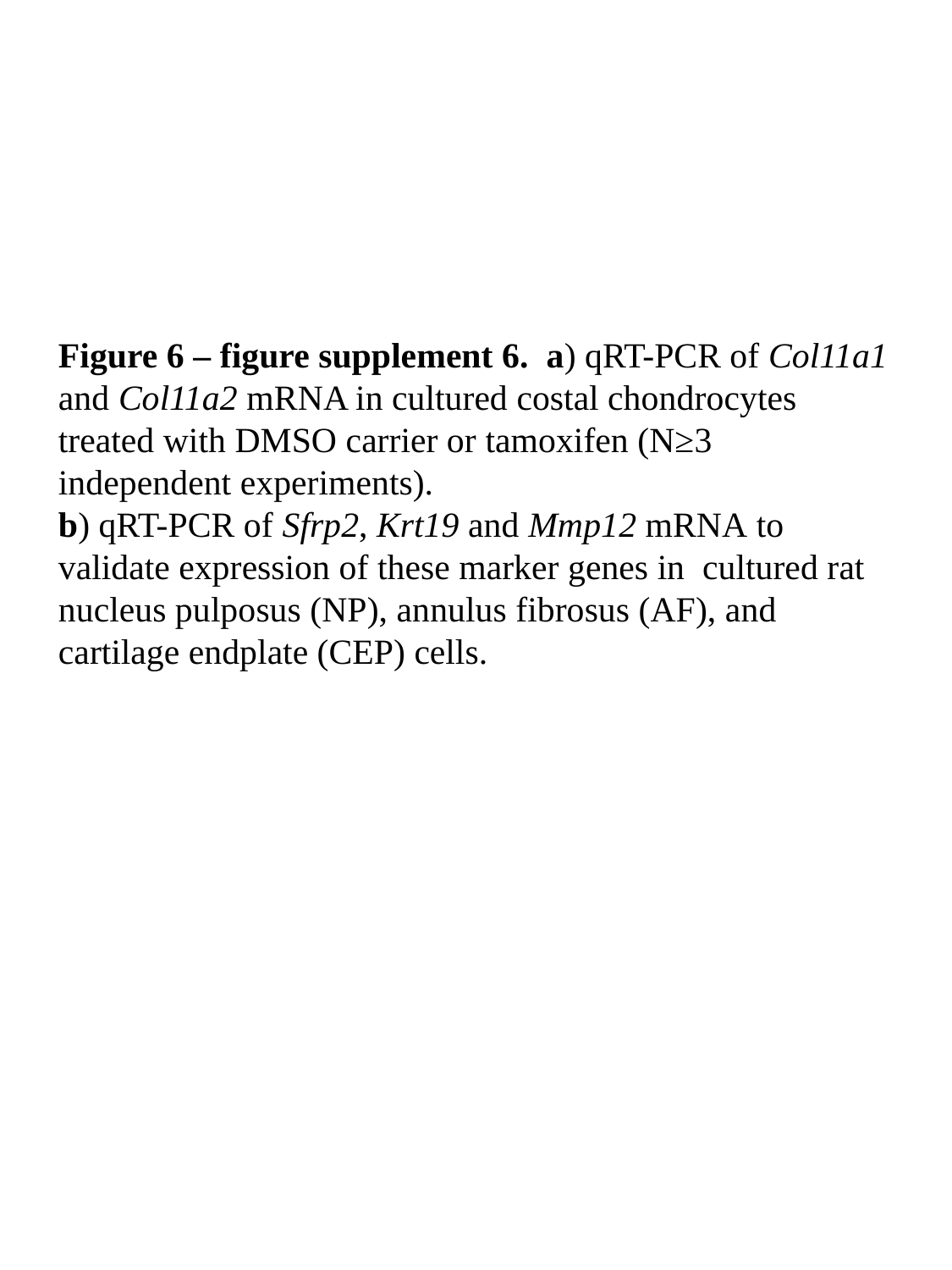

Figure 6 – figure supplement 6. a) qRT-PCR of Col11a1 and Col11a2 mRNA in cultured costal chondrocytes treated with DMSO carrier or tamoxifen (N≥3 independent experiments).
b) qRT-PCR of Sfrp2, Krt19 and Mmp12 mRNA to validate expression of these marker genes in cultured rat nucleus pulposus (NP), annulus fibrosus (AF), and cartilage endplate (CEP) cells.
